# A spatially anchored transcriptomic atlas of the human kidney papilla identifies significant immune injury and matrix remodeling in patients with stone disease

**DOI:** 10.1101/2022.06.22.497218

**Authors:** Victor Hugo Canela, William S. Bowen, Ricardo Melo Ferreira, James E. Lingeman, Angela R. Sabo, Daria Barwinska, Seth Winfree, Blue Lake, Ying-Hua Cheng, Kaice A. LaFavers, Kun Zhang, Fredric L. Coe, Elaine Worcester, Sanjay Jain, Michael T. Eadon, James C. Williams, Tarek M. El-Achkar, the Kidney Precision Medicine Project

## Abstract

Kidney stone disease causes significant morbidity and increases health care utilization. The pathogenesis of stone disease is not completely understood, due in part to the poor characterization of the cellular and molecular makeup of the kidney papilla and its alteration with disease. We deciphered the cellular and molecular niche of the human renal papilla in patients with calcium oxalate (CaOx) stone disease compared to healthy subjects using single nuclear RNA sequencing, spatial transcriptomics and high-resolution large-scale multiplexed 3D and Co-Detection by indexing (CODEX) imaging. In addition to identifying cell types important in papillary physiology, we defined subtypes of immune, stromal and principal cells enriched in the papilla, and characterized an undifferentiated epithelial cell cluster that was more prevalent in stone patients. Despite the focal nature of mineral deposition in nephrolithiasis, we uncovered a global injury signature involving multiple cell types within the papilla, characterized by immune activation, oxidative stress and extracellular matrix remodeling. The microenvironment of mineral deposition had features of an immune synapse with antigen presenting inflammatory macrophages interacting with T cells, and an immune repertoire ranging from inflammation to fibrosis. The expression of MMP7 and MMP9 was associated with stone disease and mineral deposition, respectively. MMP7 and MMP9 were significantly increased in the urine of patients with CaOx stone disease compared to non-stone formers, and their levels correlated with disease activity in stone formers. Our results define the spatial molecular landscape and specific pathways contributing to stone-mediated injury in the human papilla, and identify potential urinary biomarkers.

## Introduction

The prevalence of kidney stone disease, or nephrolithiasis, is increasing in the US and around the world. Nephrolithiasis is associated with significant morbidity, impaired quality of life and significant health care utilization. This disease is complex, with a multifactorial etiology influenced by genetic and environmental factors (Worcester and Coe 2010) (Howles and Thakker 2020). Despite decades of innovation and efforts by researchers to describe its pathophysiology, the precise mechanisms contributing to kidney stone formation remain poorly understood (Khan, Canales, and Dominguez-Gutierrez 2021). A key factor leading to this knowledge deficit is the paucity of data on the cellular and molecular makeup of the kidney papilla and its alteration with stone disease. In addition, the spatial distribution of various cell types and their association with mineral deposition (such as Randall’s plaque) during stone disease is largely unknown.

Experimental studies in rodent models of crystal formation and mineral deposition suggest that stone disease may be driven by inflammation, oxidative stress and osteogenic-like changes in the kidney papilla (Khan et al. 2012) (Joshi, Clapp, et al. 2015) (Khan et al. 2016) (Taguchi et al. 2017). However, human data in stone patients are often limited (Khan, Canales, and Dominguez-Gutierrez 2021). There is some evidence to suggest a role of the immune system in the pathogenesis of stone disease. For example, proteins associated with immune cell activation have been discovered in proteomic studies of kidney stones from patients (Mushtaq et al. 2007) (Canales et al. 2010) (Okumura et al. 2013) (Kusmartsev et al. 2016) (Tang, Nikolic-Paterson, and Lan 2019) (Xia et al. 2021). However, it remains unclear if the identified proteins are directly involved in the formation of CaOx stones or if they are simply a byproduct of non-stone related events (Witzmann et al. 2016). The enrichment of specific immune proteins can depend on the type of stone, such as the preponderance of neutrophil proteins in brushite compared to calcium oxalate (CaOx) stones (Makki et al. 2020).

Renal papilla samples from stone formers are very challenging to obtain as they require a biopsy during a nephrolithotomy surgical procedure. Molecular data from such specimens is more limited. In CaOx stone formers, Taguchi et. al. showed enriched gene expression of pro-inflammatory M1 macrophages by bulk microarray analysis of human papillary samples, thereby supporting an important role of macrophage activity in kidney stone disease (Taguchi et al. 2016) (Taguchi et al. 2021a). However, the complexity of immune cell types, their distribution and spatial neighborhoods are not known in humans. Such knowledge could be important to understand the mechanisms of stone pathogenesis, particularly to determine if immune activation is widespread or limited to cell niches associated with mineral deposition in the papilla.

Indeed, it is unclear if the underlying pathology in stone disease is limited to areas of plaque deposition or if it is more diffuse across the papilla. Technological advances such as single cell and spatial transcriptomics, and large-scale high-resolution imaging, allow for the spatial definition of cell types and states based on transcriptomics and protein markers. These technologies can advance our understanding of the pathogenesis of nephrolithiasis by defining spatial niches of various cell types/states in stone disease and address existing gaps.

In this work, we procured difficult to obtain human kidney papilla biopsy specimens from stone formers and reference nephrectomy tissues to create a spatially anchored transcriptomic atlas of the renal papilla using integrated single nuclear RNA sequencing (snRNAseq) and spatial transcriptomic (ST). By using high-resolution large-scale multiplexed 3D and Co-Detection by indexing (CODEX) imaging, we defined the spatial localization and niches of specific cell subtypes of the human papilla and the changes in this landscape with stone disease. We discovered that areas of mineral depositions are immune active zones consisting of immune injury and matrix remodeling genes that affect multiple cells types extending beyond areas surrounding mineralization. Our studies also identified MMP7 and MMP9 as potential urinary biomarkers associated with stone disease and its activity.

## Materials and Methods

### Human samples sources

#### a- Single nuclear RNA sequencing sample sources

snRNAseq data from 36 subjects comprising 203,702 nuclei were obtained from the Human BioMolecular Atlas Program (HuBMap) (https://hubmapconsortium.org/hubmap-data/) and Kidney Precision Medicine Project (KPMP) (https://www.kpmp.org) datasets that are now publicly available (GSE169285) (Lake et al. 2021). Single cells were isolated from frozen tissues as previously described (Lake et al. 2021). Of the 36, five subjects provided two samples each for a total of 41 samples. The presence of cortex, medulla, or papilla was defined by adjacent cross sectional Periodic Acid Schiff-stained histological sections. In these 41 samples, there were 29 samples with cortex, 14 samples with medulla, and five samples with papilla. Seven subjects’ samples contained both cortex and medulla. Of the five papilla samples, three were from CaOx stone formers obtained at Indiana University (see below) and two were from non-stone formers.

#### b- Human kidney papillae samples

Patients with idiopathic CaOx stones underwent a renal papillary biopsy procedure during a clinically indicated percutaneous nephrolithotomy for stone removal (Evan et al. 2003), approved by the institutional review board (IRB) of Indiana University (IRB # 1010002261). Informed consent was obtained from all study participants. Human reference (non-stone formers) nephrectomy papillary specimens without evidence of renal disease were obtained from the Biopsy Biobank Cohort of Indiana (Eadon et al. 2020) (IRB #1906572234). Following extraction, papillary tissues were frozen on dry ice in optimal cutting temperature (OCT) medium. For spatial transcriptomic and CO-Detection by indexing (CODEX) imaging studies, two reference and three CaOx papillary specimens were used. For validation studies with 3D imaging, kidney papillary sections were obtained from four additional reference and four CaOx biopsy specimens. Available clinical and demographic variables for these specimens along with the distributions of the various spatial assays performed are outlined in **Supplemental table 1**.

#### c- Cohorts for urine MMP7 and MMP9 measurements

Urine specimens from 20 healthy participants (Normal) with no personal or family history of kidney stones and 18 patients with a history of CaOx stones (Non-active stone formers) were obtained from an ongoing study at the University of Chicago (IRB protocol 09-164B). Participants were studied in the General Clinical Research Center at the University of Chicago, and the urine specimens used in these studies were obtained during the same morning period. Urine samples from 18 patients with active stone disease were obtained during elective percutaneous nephrolithotomy for stone disease at Indiana University (IRB protocol 1010002261). Demographic and relevant clinical characteristics of these subjects are presented in **Supplemental table 2**.

##### snRNAseq and processing

From an integrated HuBMAP and KPMP atlas of renal cell types (Lake et al. 2021), the snRNAseq portion of the Seurat object, including papilla samples, was reproduced to identify cell type clusters. For quality control, 10X snRNAseq cell barcodes passing 10X Cell Ranger filters were used for downstream analyses. All mitochondrial transcripts were removed. Doublets were identified and removed with DoubletDetection software (v2.4.0) (Gayoso, Shor, and Carr 2020). The 41 samples were merged and only nuclei barcodes with more than 400 and less than 7500 genes detected were maintained in the merged atlas. A gene unique molecular identifier (UMI) ratio filter was applied using Pagoda2 to remove low quality nuclei (github.com/hms-dbmi/pagoda2) (Barkas et al. 2021). Expression was calculated in 10X Cell Ranger v3 after demultiplexing and barcode processing. The GRCh38 (hg38) reference genome was used for the snRNAseq and subsequent spatial transcriptomic datasets.

##### snRNAseq clustering and annotation

snRNAseq cluster definitions were adopted from the current version of the combined HuBMAP and KPMP atlas (Lake et al. 2021). Briefly, nuclei were clustered with pagoda2. Total counts per nucleus were normalized, batch effect was corrected, and principal component analyses were performed using all significant variant genes (N= 5526). Initial cluster identities were determined by the infomap community detection algorithm. Using a primary cluster resolution of k = 100, all principal components and annotations were imported into Seurat v4.0.0 to create a merged uniform manifold approximation and projection (UMAP). Standardized anatomical and cell type nomenclature was used to annotate cell types and subtypes, based on the collaborative KPMP and HuBMAP definitions. These definitions were based on published datasets and the expertise of consortium pathologists, biologists, nephrologists and ontologists (El-Achkar et al. 2021) (Lake et al. 2019) (Gerhardt et al. 2021) (Chen, Chou, and Knepper 2021) (Ransick et al. 2019) (Borner et al. 2021). Specifics of the cluster decision tree algorithm have been previously described (Lake et al. 2021). Within the integrated HuBMAP and KPMP atlas, putative adaptive and degenerative cell states were identified in epithelial sub-clusters with at least one of the following: reduced genes detected, higher mitochondrial transcript amount, higher ER associated transcript number, increased expression of known injury markers (e.g., IGFBP7, HAVCR1, LCN2, CST3, etc.), or enrichment in samples with acute or chronic kidney disease. Markers of these cell states were identified using the Seurat function “FindConservedMarkers” with the following parameter settings: grouping.var = “condition.l1”, min.pct = 0.25, and max.cells.per.ident = 300. The gene set for the adaptive and degenerative cell states were trimmed to include only enriched genes at a p value < 0.05 and mean log2 fold change > 0.6. Due to the consistent loss of epithelial cell type specific markers, all nuclei within the adaptive and degenerative cell state clusters were merged into a single undifferentiated epithelial cluster within the papilla snRNAseq UMAP.

##### Visium Slide preparation, mRNA extraction and sequencing

Frozen 10 µm sections were mounted onto etched frames of the Visium spatial gene expression (VSGE) slides according to 10x Genomics protocols (*Visium Spatial Protocols—Tissue Preparation Guide*, Document Number CD=G000240 Rev A, 10x Genomics). Tissue sections were fixed with methanol, subsequently stained with hematoxylin and eosin (H&E) and imaged by bright-field microscopy. Microscopic images were acquired according to protocols described in 10x Genomics protocols (Technical Note - *Visium Spatial Gene Expression Imaging Guidelines*, Document Number CG000241, 10x Genomics). H&E-stained sections were imaged with a Keyence BZ-X810 microscope equipped with a Nikon 10× CFI Plan Fluor objective at 0.7547 um/pixel and image resolution of 1920×1440. Images were collected as mosaics of 10x fields. Stained tissues were permeabilized for 12 minutes. mRNA bound to oligonucleotides on the capture areas of the Visium slides was extracted. cDNA libraries were prepared with second strand synthesis and sequenced utilizing the NovaSeq 6000 Sequencing system (Illumina) in the 28 bp + 120 bp paired-end sequencing mode. Post sequencing quality metrics are included in **Supplemental table 1**.

##### Spatial transcriptomics expression analysis

Using Space Ranger 1.2.0 with the reference genome GRCh3-2020-A, samples were mapped, and counts were generated that corresponded to the barcoded 55 μm spot coordinates within each fiducial frame. This allowed association of read counts with their location within the H&E image. Space Ranger calculated differential expression between assigned clusters using sSeq and edgeR. The data were normalized by SCTransform and merged to build a unified UMAP and dataset as previously described (Melo Ferreira, Freije, and Eadon 2021). All feature plots show expression after normalization.

##### Selection of histologic phenotypes

Kidney stone papillary tissue biopsies were manually annotated utilizing the 10x Genomics Loupe Browser (10x Genomics Loupe Browser 5.0.0). Specific regions of the stone tissue papilla were categorized as either non-mineralized, contiguous to mineral, or mineralized regions. Spots directly overlying evident mineral precipitation were designated as mineralized; spots falling within a three-spot radius of the annotated mineral precipitation were categorized as “contiguous” to mineral; all remaining spots were designated as non-mineralized.

##### Pathways and gene expression analysis

Differential expression comparisons across samples were performed using R and the R packages ReactomePA and ClusterProfiler as previously described by Ferreira et al. (Melo Ferreira et al. 2021). The DEGs in each comparison were found with the Seurat function FindMarkers and tested with a Wilcoxon’s rank sum test. Pathway enrichment for those genes was performed with the R packages ReactomePA (Yu and He 2016) and ClusterProfiler (Yu et al. 2012).

##### Label Transfer

Seurat v3.2 was used to accomplish label transfer from snRNA-Seq cell types and cell states to spatial transcriptomic spots in each sample. For deconvolution analyses, a Seurat v3.2 anchor methodology was used to transfer single-cell cluster information to Visium data as described (Melo Ferreira et al. 2021). Each spot receives a probability or transfer score for its association with a given snRNA-Seq cell type or cell state cluster. The transfer scores are summed, and each spot is deconvoluted with the fractions of the spot corresponding to the relative proportion of transfer score of each contributing snRNAseq cluster. A pie chart is displayed over the spatial transcriptomic sample image.

##### Tissue Processing, Immunofluorescence Staining and Large-Scale 3D Confocal Imaging

Papillary biopsies were immediately immersed in OCT medium and frozen on dry ice. For immunofluorescence analysis, tissues were cryosectioned (Leica Biosystems, Wetzlar, Germany) at 20 μm thick sections. Sections were washed in phosphate-buffered saline (PBS), fixed for 4 hours at room temperature (RT) in 4% paraformaldehyde. Next, the sections were washed in PBS two more times, and then blocked in 10% normal donkey serum for two hours at RT. Primary antibodies were added in blocking buffer and tissues were incubated at RT overnight. When targeting intracellular antigens, permeabilization was performed using 0.2% Triton X (Santa Cruz Biotechnology, Inc., Dallas, TX) (Gildea, 2017) (El-Achkar et al. 2007). The following primary antibodies were used for detection: anti-aquaporin 1 (Santa Cruz Biotechnology, Inc., Dallas, TX; sc-9878), anti-CD68 (Agilent Technologies, Santa Clara, CA; M087601), anti-phospho-c-JUN (Cell Signaling Technology, Danvers, MA; 9261). After washing with PBS, the following Alexa Fluor (ThermoFisher Scientific, Waltham, MA) dye-conjugated secondary antibodies were added: donkey anti-mouse-488, anti-mouse-568 and anti-rabbit-647. 4′,6-Diamidino-2-phenylindole (DAPI) (Abcam, Cambridge, United Kingdom; ab228549) was used for staining nuclei. Subsequently, sections were washed three times for 30 minutes each in PBS and then fixed in 4% paraformaldehyde for an additional 15 minutes. After a final wash in PBS for 30 minutes, sections were mounted on a glass slide using ProLongTM Glass Antifade Mountant, (Thermo Fisher Scientific, Waltham, MA; REF# P36980). Images were sequentially acquired in 4 separate channels using the Leica SP8 confocal microscope and collecting whole volume stacks using 20xNA 0.75 objective with 1.0-μm spacing. Stacks were stitched using Leica LAS X software to generate large-scale 3D images. A negative control without primary antibody was used to ensure the absence of nonspecific binding of secondary antibodies. Microscope settings were identical among imaging sessions for each specimen.

##### 3D Tissue Cytometry

3D tissue cytometry was performed on image volumes using the volumetric tissue exploration and analysis (VTEA) software (Winfree et al. 2017). Segmentation settings were adjusted to yield the best result which was verified visually by sampling random fields within each image stack. Fluorescence from phospho-c-Jun and CD68 was associated with nuclei by 3D morphology and displayed on a scatterplot as individual points, allowing gating of specific cell populations based on fluorescence intensities. Results were reported as percentage of total cells segmented in each large-scale image volume (Makki et al. 2020).

##### CODEX Antibody Conjugation and Validation

A total of 32 antibodies were used for CODEX and 19 of them were conjugated in-house using a protocol outlined by Akoya Biosciences (Akoya Biosciences, Inc. CODEX® User Manual, Menlo Park, CA). Conjugation of antibodies to their assigned barcodes, they were first reduced using a “Reduction Master Mix” (Akoya Biosciences, Menlo Park, CA). Lyophilized barcodes were then resuspended using the Molecular Biology Grade Water and Conjugation Solution. The barcode solution was then added to the appropriate reduced antibody and incubated for two hours at RT. After incubation, the newly conjugated antibody-barcode was purified in a three-step wash/spin process and stored at 4°C. Successful conjugation was validated via gel electrophoresis as well as immunofluorescent staining and confocal imaging.

##### CODEX Imaging

Human renal tissue sections of 10 µm were cut from OCT blocks onto poly-L-lysine coated coverslips. Sections were prepared as detailed by the manufacturer’s instructions (Akoya Biosciences, Menlo Park, CA). Tissue retrieval was conducted with a three-step hydration process, followed by a PFA fixation. During fixation, an antibody cocktail of the 32 antibodies listed in **Supplemental Table 3** was made and then dispensed onto the coverslip. Tissues were allowed to incubate with the staining solution overnight at 4°C. The following day, the staining solution was washed from the tissues and a multi-step fixation occurred (Schurch et al. 2020).

The imaging of tissues was conducted at 20x resolution using the CODEX system from Akoya Biosciences and a Keyence BZ-X810 slide scanning microscope (Keyence Corporation, Itasca, IL). The resulting images were processed using the CODEX processing software (Akoya Biosciences, Menlo Park, CA) and visualized using FIJI/ImageJ.

##### Unsupervised analysis, clustering and mapping of cell types

CODEX images were segmented and analyzed using VTEA version 1.0.3, (Winfree et al. 2022). Cell clusters were identified using unsupervised Ward clustering and dimensionality reduction visualization using *t*-distributed stochastic neighborhood embedding (*t*-SNE). The identity of the cell clusters was verified by plotting their mean intensities for specific makers and directly mapping on the image volumes using nuclear overlays.

##### ELISA and Urine Collection

Urine samples were collected from patients with a history of CaOx stone formation (non-active stone formers), from CaOx stone patients immediately before undergoing surgery for stone removal (active stone formers) or from non-stone formers as described above. The urine samples were immediately frozen at -80°C. At the time of analysis, samples were thawed and cleared by centrifugation for 5 minutes at 1000xg and assayed for urine MMP7 and MMP9 by ELISA according to manufacturer’s instructions (R&D Systems, DMP700/DMP900, Minneapolis, MN). Urine creatinine was used to normalize MMP7 and MMP9 levels between non-active, active, and non-stone formers.

##### Statistics

Statistics were used within transcriptomics analyses are described above. Student’s *t*-test was used to compare the phospho-c-JUN and CD68 positive cells between patient samples and reference tissues. Urine levels of MMP7 and MMP9 were compared using ANOVA with the Tukey-Kramer post hoc test.

##### Data Availability

Raw snRNAseq data generated as part of the KPMP has been accessed from atlas.kpmp.org and from the HuBMAP at portal.hubmapconsortium.org. Visium spatial transcriptomic data is available in GEO as GSE206306.

## Results

### Identification and localization of cell types in the kidney papilla

We recently reported a comprehensive single nuclear RNA sequencing (snRNAseq) profile (over 200,000 cells) of the adult human kidney from the HuBMAP and KPMP consortia (**Figure 1A**) (Lake et al. 2021). We leveraged this dataset to gain a deeper understanding of cell type diversity in the human kidney papillae and unique features compared to the cortical-medullary cell types. Renal papilla samples contributed 20,338 nuclei to this dataset (**Figure 1B**). These samples were acquired from papillary biopsies of CaOx stone formers or were dissected from reference nephrectomies. In addition to the expected papillary cell types such as principal and intercalated collecting duct cells, descending and ascending thin limbs, papillary surface epithelial, stromal and endothelial cells (**Figure 1C**), notable differences in gene expression were observed between papillary and cortico-medullary principal (PC) and intercalated (IC) cells (**Figure 1D**). For example, the urea transporter UT2 (*SLC14A2*) and Aquaporin 2 (*AQP2)* were both upregulated in papillary PCs and ICs. In contrast, the expression of the epithelial sodium channel ENaC (*SCNN1G, SCNN1B*) and PTH receptor (*PTH2R*) were more abundant in the cortico-medullary nuclei. Further, the gene *RALYL*, encoding the RNA-Binding Raly-Like Protein and responsible for the cystic Bardet-Biedl Syndrome 1, was predominantly expressed in PCs outside the papilla (**Figure 1D**). Distinct sodium bicarbonate transporters were also expressed in papillary as compared to cortico-medullary cells. These findings highlight the unique physiological role of these papillary cells in regulating water and urea transport and acid base balance in this unique environment and may have relevance to the pathogenesis of cystic disease.

**Figure 1:**
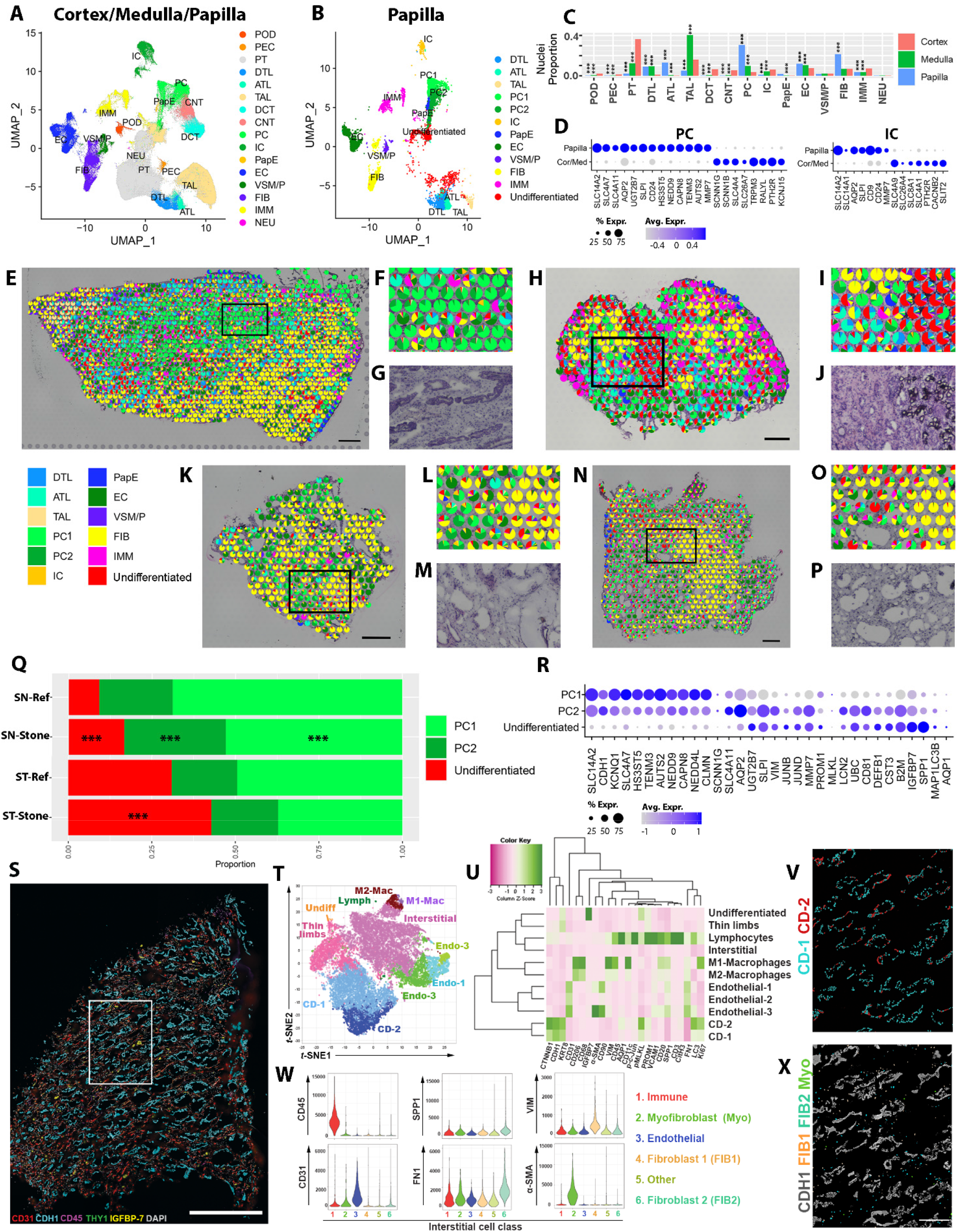
Spatially anchored cellular and molecular characterization of the human kidney papilla. (A) An integrated snRNA sequencing atlas from the KPMP and HuBMAP consortia presented as a Uniform Manifold Approximation and Projection (Umap) combining nuclei from the renal cortex, medulla and papilla. (B) A subset of nuclei specific to the papilla alone. (C) The proportion of cell type representation among nuclei in each kidney region. (D) Both papillary principal (PC) and intercalated (IC) cells had increased expression of the urate 2 transporter (*SLC14A2*) and other differentially expressed genes (DEGs) as compared to their counterparts in the renal cortices and medullae. (E) Label transfer and mapping of the snRNAseq cell classes onto spatial transcriptomic spots within a reference papilla tissue. (F) An enlarged area denoted by the box in (E) showing principal cells mapping on histologically identified collecting ducts in (G). (H) Label transfer and mapping of cell classes onto ST spots in a papilla of a stone former. (I) An enlarged area from (H) showing the undifferentiated cell signature localizing to areas of mineralization (J). (K-P) ST of additional stone papilla specimens showing the signatures of fibroblasts and PC2 (L and M) and undifferentiated cells (O and P). (Q) distribution of PC1, PC2 and undifferentiated cells between reference and stone samples across the snRNAseq and ST data. PC2 and undifferentiated cells were relatively more abundant in stone samples. (R) Gene expression signatures of PC1, PC2 and undifferentiated cells, showing a spectrum of injury in the PC2 and undifferentiated cell types. (S) Co-detection by Indexing (CODEX) multiplex imaging of a reference papilla tissue sample. (T) Unsupervised clustering and dimensionality reduction in a t-stochastic neighborhood embedding (t-SNE) plot showing various cells classes, which were validated by the level of fluorescence intensity (U) and mapping back on the image. (V) localization of CD-1 and CD-2. CD-2 cells express higher levels of injury markers (U) but are not segregated into separate tubules. (W) Re-clustering of interstitial cells based on specific markers identifies populations of fibroblasts (2 subtypes, FIB1 and FIB2) and myofibroblasts (Myo), which were mapped into the interstitium around the epithelium (X). *** denote P<0.01. Scale bars: (0.5 mm in E and K, 0.25 mm in H and N, 1 mm in S and 100µm in X)

We then sought to orthogonally validate these cell types and spatially resolve them in papillary tissue from controls and nephrolithotomies of subjects with CaOx stones. We used snRNA-seq labels to resolve the spatial transcriptomics (ST) gene expression profiles obtained from 10X Visium. (**Figure 1E-P**). The integrated analysis showed the mapping of the appropriate histological structures with the papillary tubules (**Figure 1F-G**). The majority of papillary surface epithelial cells (PapE) mapped to the outer edge as expected (**Figure 1E, H, N**). Fibroblast and immune signatures were identified in all the samples tested. We found a unique population of undifferentiated cells that were enriched in injury associated genes and lacked a transcriptomic signature specific to a unique cell identity (**Figure 1B**). Undifferentiated cells were more abundant in stone disease samples (**Figure 1Q**). ST data showed that these cells frequently localized near areas of mineralization and are likely a result of localized injury to the adjoining tubules (**Figure 1H-J**).

Further characterization of the principal cells (PC) showed that in addition to the healthy PC1 population, a PC2 population enriched with stress/injury genes was identified. These principal cell subtypes were compared to the undifferentiated epithelial cell type that also exhibited an injury signature (**Figure 1B**). Stone samples had a higher proportion of PC2 (compared to PC1) and undifferentiated cells by both snRNAseq and ST (**Figure 1Q**). Both PC1 and PC2 cells expressed canonical papillary PC markers; however, PC2 cells also expressed injury markers such as *VIM, JUNB, JUND, LCN2* and *MAP1LC3B* (LC3) (**Figure 1R**). The undifferentiated epithelial cells had greater expression of injury markers such as *PROM1, IGFBP7* and *SPP1* and likely represents a degenerative state.

To extend the transcriptomic cell type prediction to the protein level, we performed highly multiplexed CODEX imaging of reference tissues using 32 cell markers (**Figure 1S-X**). Applying unsupervised analysis and classification based on the markers used, the presence of various expected cell types was confirmed, mapped within the tissue, and aligned with the cell types from the transcriptomic datasets (**Figure 1T** and **1U**). We confirmed the presence of a significant resident immune cell population, with a predominance of macrophages with high expression of CD206 (M2-Macrophages). CODEX analyses uncovered two populations of collecting duct cells in the papilla, CD1 and CD2, which predominantly consist of principal cells based on the distribution outlines in **Figure 1C**. These two populations were differentiated by the expression of injury and renewal markers in CD2 such as LC3, p-MLKL, PROM1 and Ki67 (**Supplemental Figure 1**) and consistent with the transcriptomic injury signature uncovered in PC2 cells. We also uncovered a population of undifferentiated epithelial cells (**Figure 1T, 1U and Supplemental Figure 2**) whose protein expression profile also overlaps with the undifferentiated population uncovered by ST (high expression of IGFBP7 and PROM1). Thus, using snRNAseq, ST and CODEX we validated the cellular diversity in the papilla and identified new cell populations with plausible biological significance (see below).

Using CODEX, we then explored the population of stromal cells localized in the papillary interstitium, which was divided into separate cell classes based on an unsupervised analysis (**Figure 1W**). We uncovered and spatially mapped two populations of fibroblasts based on the expression of fibronectin (FN1), vimentin (VIM) and osteopontin (SPP1). We also identified a population of myofibroblasts with high expression of α-SMA (**Figure 1X**). The presence of fibroblasts in the papilla is consistent with our snRNAseq and ST findings in **Figure 1E-1P**.

### Transcriptomic signatures of CaOx stone formation within the renal papilla

Using the snRNAseq data, we determined the differentially expressed genes (DEGs) between CaOx stone and reference samples for each cell type in the papilla (**Figure 2 A-D, Supplemental Figure 3**). Multiple cell types in stone samples expressed increased levels of injury and stress genes such as *SPP1, CLU, LCN2, S100A11, MMP7* and *CD74*. These injury genes were similarly upregulated in the cell signatures and global gene expression in the ST stone samples (**Figure 2E-F** and **Supplemental Figure 4**). Pathway analysis detected significant enrichment of common pathways related to protein translation, among the cells from patients with stone disease. We also observed enrichment of pathways of interest (**Figure 2 A-D**), such as: leukocyte activation, response to oxidative stress, ossification, extracellular matrix organization in most cell types, which overlapped with enriched pathways obtained independently from the spatial transcriptomic analysis (**Supplemental Figures 3 and 4**). The concordant results between technologies increase the confidence in the biological relevance of these pathways in the pathogenesis of stone disease.

**Figure 2.**
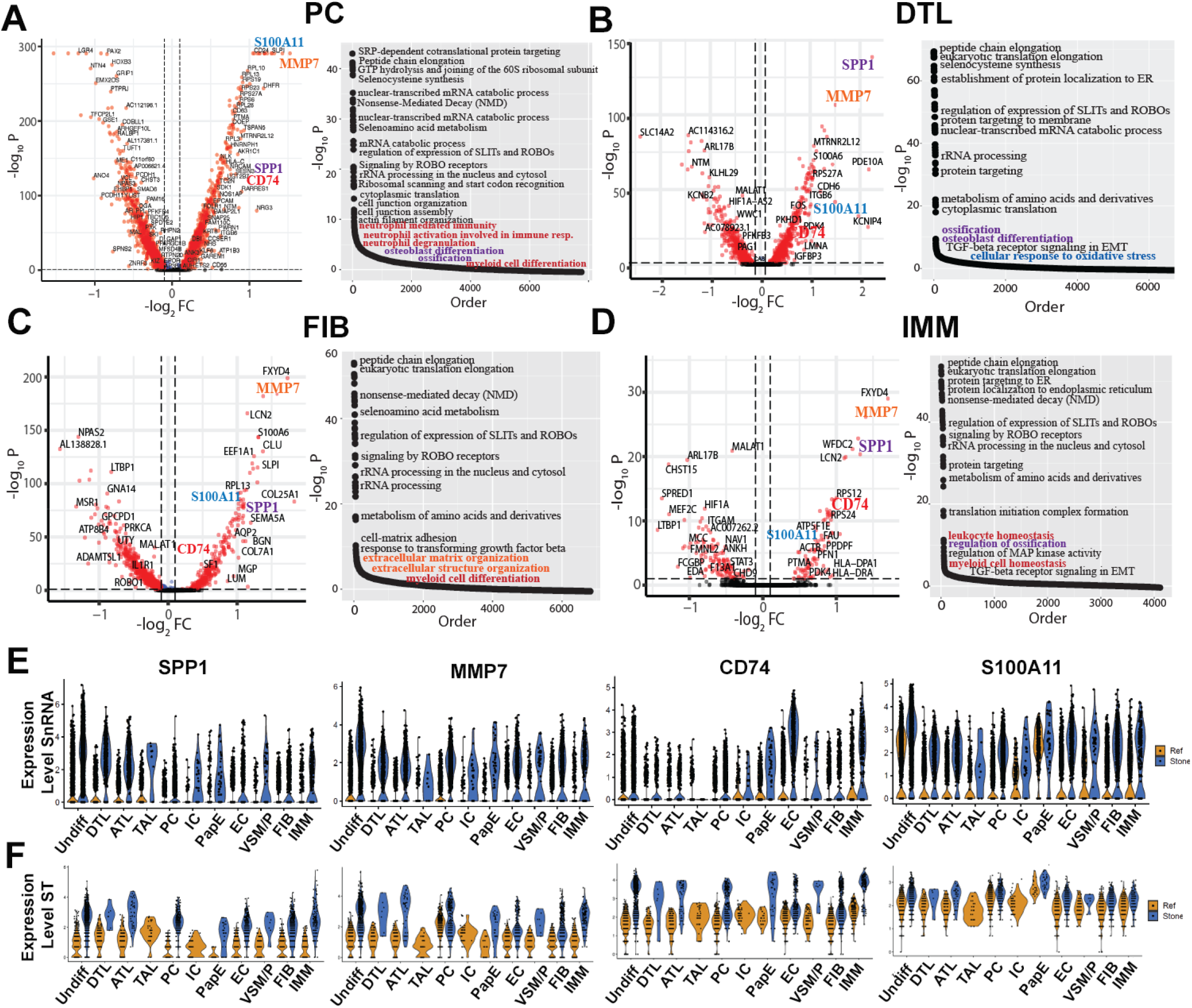
Differentially expressed genes (DEGs) induced by stone disease in various cells within the human papilla. (A-D) DEGs and enriched pathways in papillary cells by snRNAseq, showing few examples: principal cells (PC), descending thin limbs (DTL), fibroblasts (FIB) and immune cells (IMM). CaOx stone disease induces similar increases in gene expression in various papillary kidney cells. Pathway analysis of snRNA expression detected significant upregulation of ossification (purple text), extracellular matrix organization (orange text), response to oxidative stress (blue text) and leukocyte activation pathways (red text). (E) Violin plots showing differential expression in reference and stone specimens across all papillary cell types of major genes consistently induced by stone disease. (F) Expression levels of the genes shown in (E) in the cell signatures mapped on the spatial transcriptomics data in the refence and stone samples.

To further gain insights into the pathogenesis of injury in stone disease, we looked in more detail *MMP7* expression, a gene involved in injury and matrix remodeling and consistently upregulated in the papillae from patients with stone disease. **Figure 3** shows the spatial and relative gene expression maps of *MMP7*. Interestingly, in reference tissue, *MMP7* was confined to CDs, but in stone patients *MMP7* showed a diffuse expression pattern, which was consistent with the generalized increased expression observed in most cell types from the snRNAseq and combined analysis (**Figure 2E** and **2F**).

**Figure 3.**
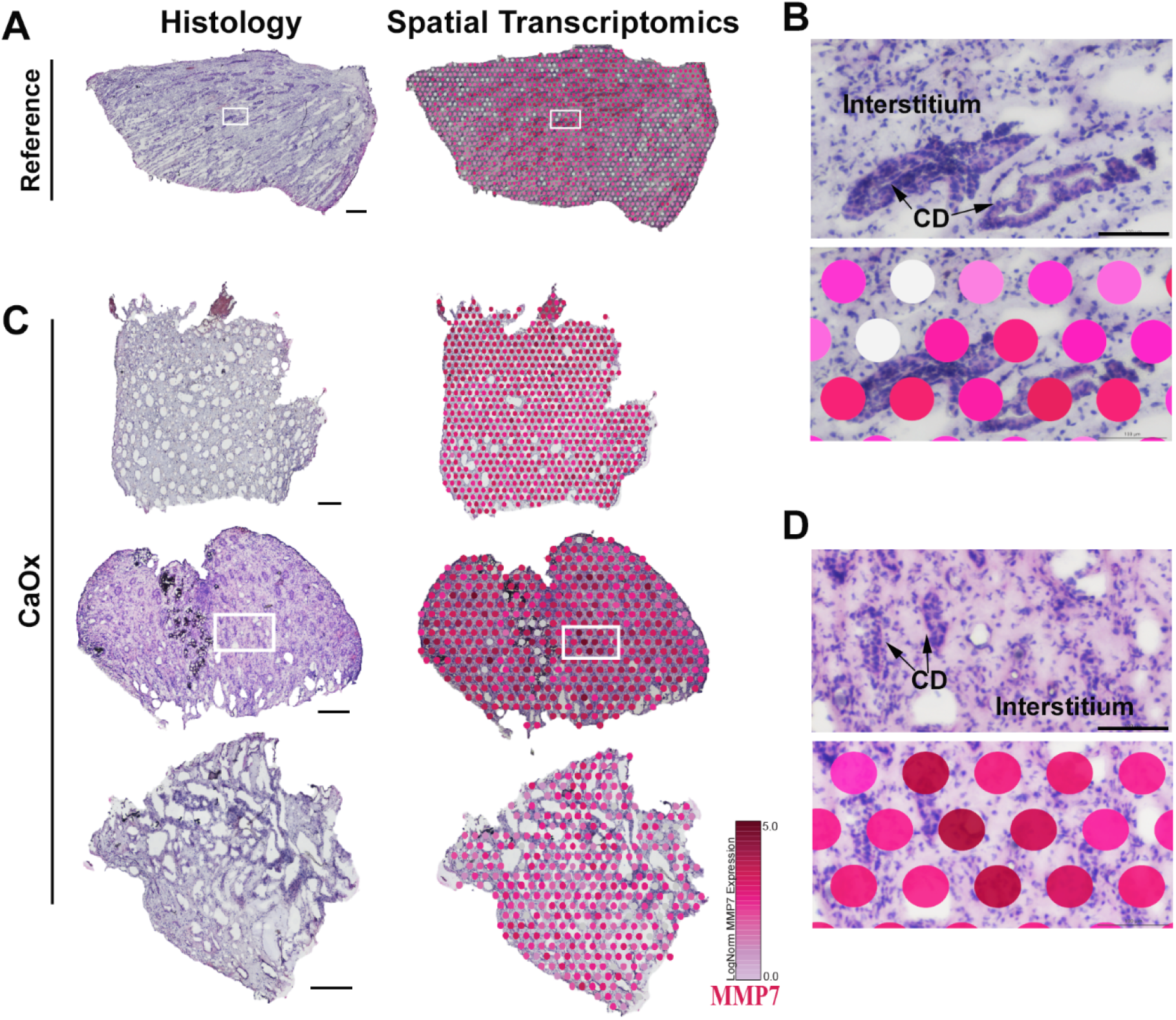
MMP7 expression in the papilla. Spatial transcriptomics analysis comparing control (Reference) and three different CaOx stone patient biopsies. In reference tissue, MMP7 expression is localized to collecting ducts. In stone disease, MMP7 expression is diffusely increased and encompasses various papillary cells and structures, which is consistent with the snRNAseq expression and the expression signature mapping on ST shown in Figure 2E and F. Scales bars in (A) and (C): 0.5 mm for reference and top stone sample, 0.25 mm for other 2 specimen; for (B) and (D): 0.1 mm.

### Regional analysis of mineralized and non-mineralized tissues

Mineral deposition such as Randall’s plaque formation is thought to play an important role in the formation of stone disease. However, plaque formation is frequently focal, and the cellular and molecular alterations present in the microenvironment of mineral deposits could be important in the pathogenesis of this disease. Supervised analysis of spatial transcriptomic profiles based on the selection of mineral deposits in a stone forming papilla are shown in **Figure 4**. The results indicate that transcript signatures of genes such as *NEAT1, CHIT1, LYZ, SPP1* and *MMP9* are upregulated in areas contiguous to mineralized tissue compared to regions more distant (non-contiguous) to mineral (**Figure 4**). These genes are also associated with pathways of leukocyte activation and response to oxidative stress (**Supplemental Figure 5**). The expression of genes involved in macrophage activation such as *MMP9* and *CHIT1* (**Figure 4B**) was upregulated in the area of mineralization as compared to the non-contiguous region.

**Figure 4.**
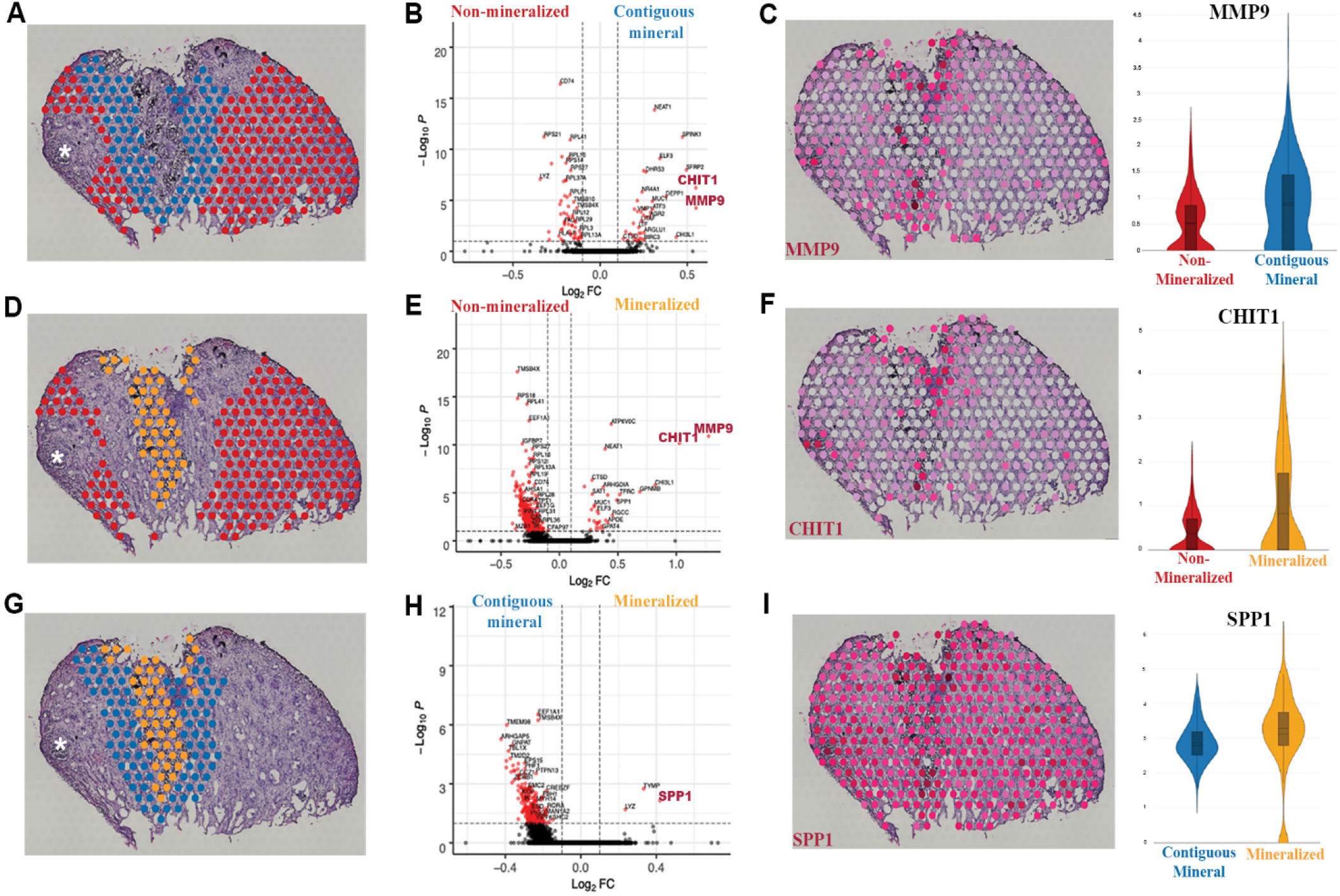
Signatures of injury and inflammation are localized to regions of mineralization in the kidney papilla from stone disease. (A) Regions of non-mineralization compared to areas contiguous to mineral display differentially expressed genes (DEGs) associated with pathways leukocyte activation such as MMP9 and CHIT1 (B). MMP9 expression localizes in areas contiguous to mineral and in regions of mineralization (C). Comparisons between areas of non-mineralization and areas of mineralization (D) also display DEGs such as MMP9 and CHIT1 (E). CHIT1 expression also appears to be more robust in areas of mineralization (F). Regional comparison between areas contiguous to mineral and areas of mineralization is shown in (G). SPP1, TYMP and LYZ were differentially expressed in areas of mineralization in this CaOx biopsy specimen (H). SPP1 appears to be localized to regions of mineralization but also displays relatively high expression throughout this stone forming papilla (I).Violin plots show Log-Normalized values. Asterisk in A, D and G denote that the analysis excluded the areas of mineralized plug and was restricted to Randall’s plaque.

We next wanted to better define the niche of the mineralization nidus and understand its contribution to injury, particularly with our findings of enrichment of immune cell marker genes and their association with mineral deposition. We used CODEX multiplexed imaging in conjunction with a panel of antibodies directed to different immune cell types (**Figure 5**). We took advantage of the autofluorescence properties of mineral deposits (**Figure 5A-B**), whereby the plaque can be identified without staining (Makki et al. 2020) (Winfree et al. 2021). The main cell clusters identified in refences tissue (**Figure 1T-U**) were also identified here, except that there was marked expansion of the immune clusters (**Figure 5B)**. Additional unsupervised analysis (**Figure 5C**) was performed on the CD45+ cell clusters from the initial analysis (**Figure 5B**), which resolved immune cells into major lymphocyte and myeloid clusters. The latter can be broadly divided into three categories based on markers such as CD68 and CD206, which correspond to an inflammatory (M1), alternatively activated (M2) and intermediate phenotype (M1/M2). Mapping back immune cells to the tissue images reveals that certain areas of plaque deposition serve as a nidus for immune activation (**Figure 5D**), where the deposited mineral is surrounded by inflammatory macrophages (M1 and M1/M2), which are known to be antigen presenting cells (APCs) (**Figure 5E**). At the periphery of this immune nidus, these APCs interact with CD4+ T and B cells, a phenomenon consistent with classic immune activation and synapse formation (**Figure 5E**). Interestingly, not all the plaque areas are involved with immune activation, and neighboring areas with mineral did not have significant immune cells. We then examined the distribution of other interstitial cell populations such as fibroblasts and myofibroblast in these areas (**Figure 5F-G**). The plaque areas without immune activity had significant fibroblast infiltration (**Figure 5H**). Cumulatively, these data demonstrate that the presence of plaque is immunogenic. It elicits a varied immune response with a defined progression from myeloid to a mixed lymphocytic infiltrate and a non-overlapping fibrogenic response consisting of activated fibroblasts and myofibroblasts for repairing the injured tissue.

**Figure 5.**
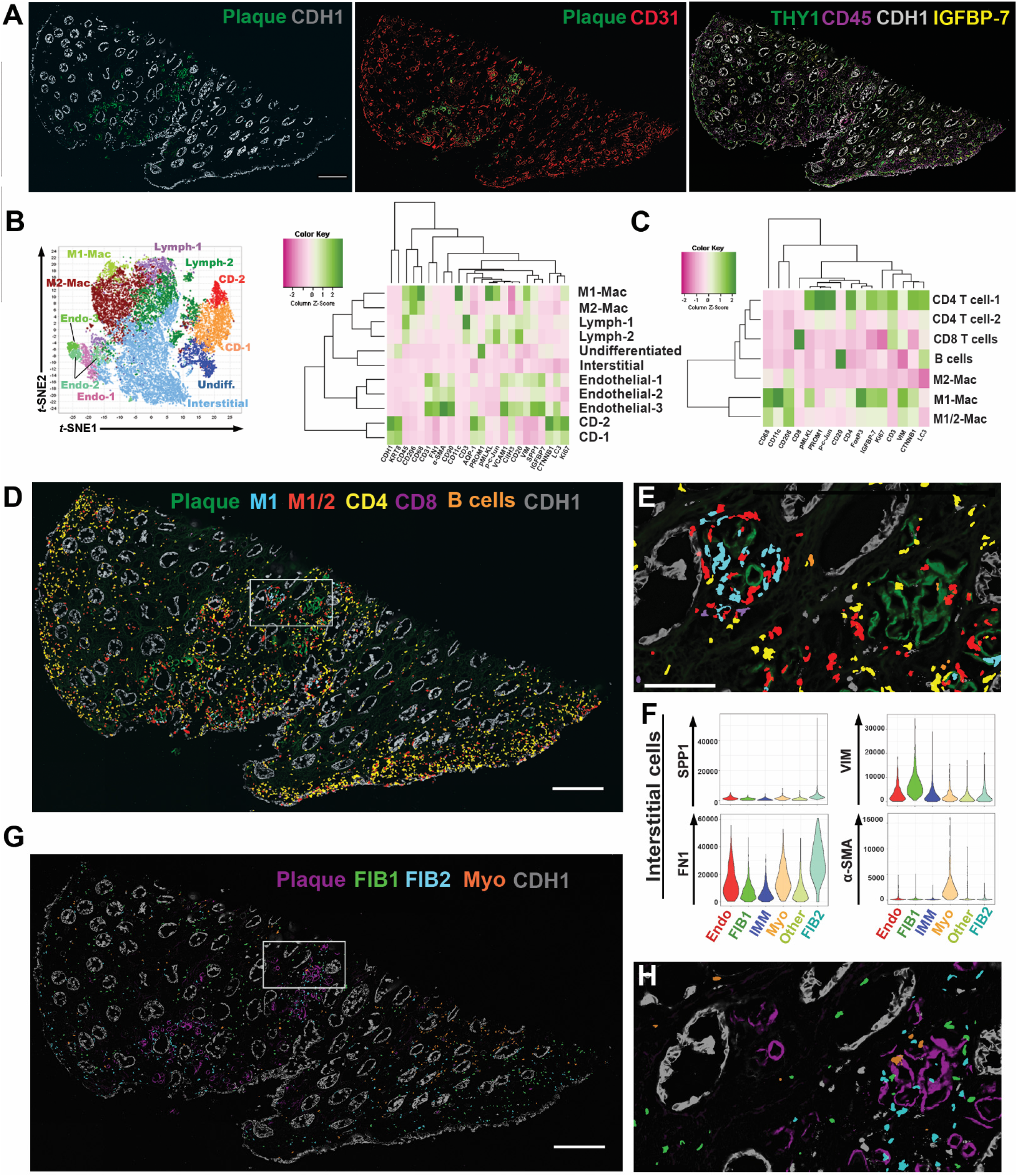
CODEX imaging of a papilla with mineral deposition identifies various stages of immune activation and fibrosis around the plaque. (A) CODEX imaging showing a unique autofluorescence of the plaque, which can be easily delineated from the epithelial and vascular cells. (B) Unsupervised analysis and clustering identify similar clusters as in the reference specimen (Figure 1), with extensive expansion of the immune clusters. (C) Re-clustering and analysis of the immune cells identifies all the major subtypes of leukocytes. Macrophages had an intermediate phenotype between M1 and M2 based on the co-expression of specific markers. (D) Mapping of immune cells in the tissue reveals niches of immune activity in certain plaque areas (area of mineral on left in (E), which is a high magnification view of the boxed area in (D)) with features of antigen presentation (interaction of antigen presenting macrophages with T and B cells) and a diffuse activated T cells response, particularly towards the papillary endothelium. Interstitial cells (F), particularly fibroblasts and myofibroblasts, were abundant throughout the tissue (G), but fibroblasts were concentrated in certain areas of mineral deposition with reduced immune activity (area of mineral on right in (H), which is a high magnification view of the same boxed area in (D) and (G)), suggesting progression from inflammation to fibrosis in neighboring areas within the same tissue. Scales bars: 500µm in (A), (D) and (G); 100 µm for (E).

### Large scale 3D imaging and tissue cytometry establishes inflammatory stress signaling and macrophage activation as cardinal features of stone disease

Next, we wanted to determine if immune activation and oxidative injury are cardinal features in the papilla of CaOx stone forming patients, independent of large mineral deposits. Using quantitative large-scale 3D imaging and tissue cytometry on papillary tissue specimens from controls and subjects with CaOx stone disease and no visible mineral deposits (N=4 each group), we quantified the expression and distribution of phosphorylated c-JUN (p-c-JUN, marker of oxidative stress and stress kinase activation) and CD68 (marker for inflammatory macrophages) (**Figure 6**). Patients with CaOx stones compared to controls had a significantly higher abundance of CD68+ macrophages (3.4 +/-1.4 vs. 1.2 +/-1.1 % of total cells, respectively; p=0.03) and p-c-JUN+ cells (12.4 +/-4.0 vs 4.0 +/-3.8 % of total cells, respectively, p=0.01) than non-stone formers. Macrophage infiltration was diffuse, and activation of c-JUN was not restricted to a specific cell type (**Figure 6D**), which is consistent with the findings from snRNAseq and transcriptomic analyses suggesting increased stress signaling across multiple cell types in the papillae of stone patients.

**Figure 6.**
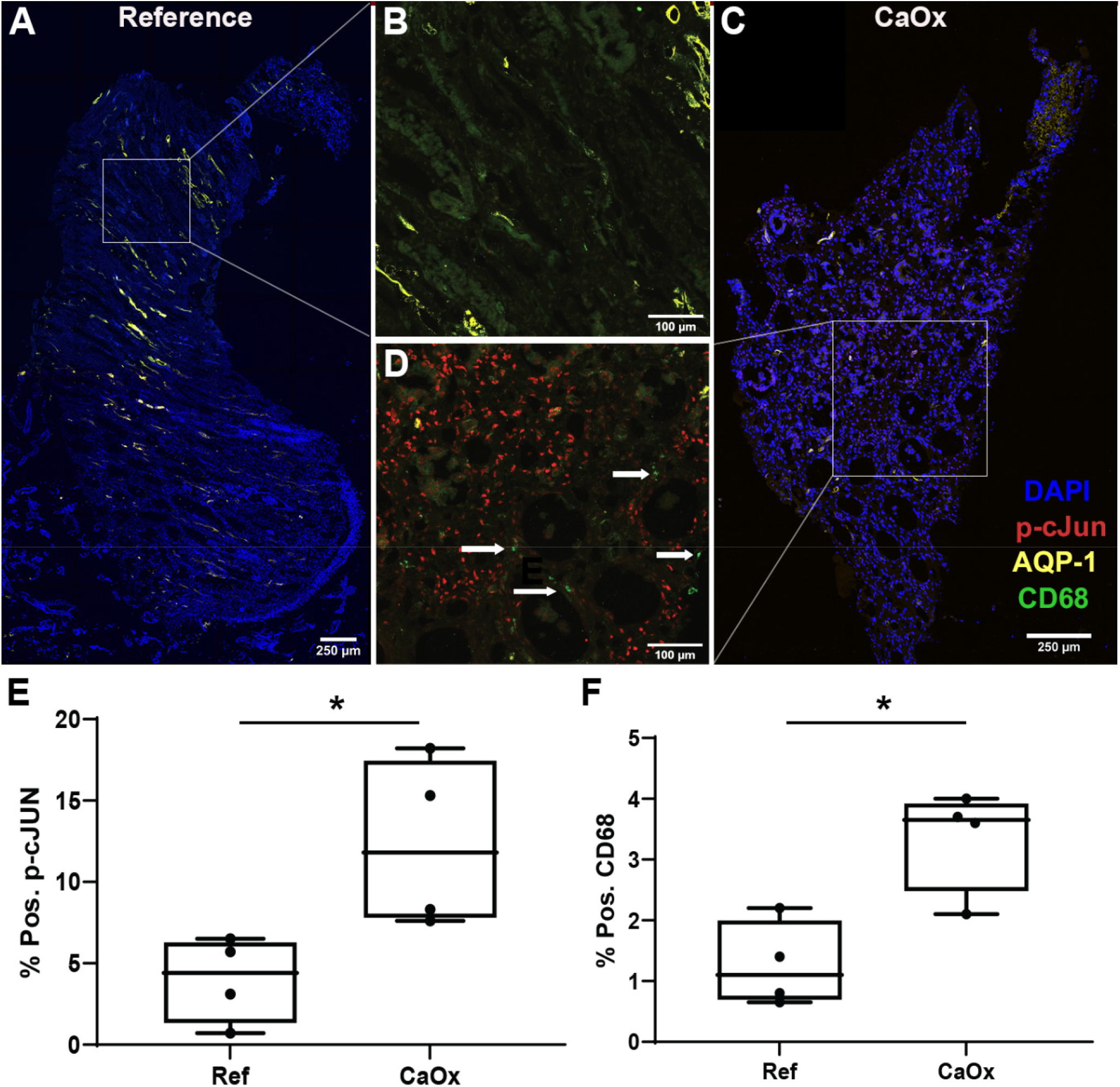
Protein markers of oxidative stress (ROS) and macrophage activation are diffusely increased in biopsies of stone patients. (A-D) Representative multi-fluorescence confocal images of kidney papillary biopsies from stone patients and reference nephrectomies tissue specimens (N=4 per group) stained for phospho-c-JUN (p-c-Jun, marker of ROS), CD68 (activated macrophages) and Aquaporin1 (AQP-1, marker for thin descending limbs and descending vasa recta). Images were analyzed using volumetric tissue exploration and analysis (VTEA) software and the resulting outcomes are shown in (E) and (F) for p-c-JUN and CD68 as percentages of total cells in each tissue. Boxed areas in (A) and (C) are enlarged in (B) and (D), respectively.

### MMP7 and MMP9 levels are increased in urine of kidney stone patients and correlate with disease activity

The snRNAseq and ST datasets revealed upregulation of *MMP7* gene expression in the papillae of patients with stone disease and the upregulation of *MMP9* gene expression in regions of mineralization. While these 2 matrix remodeling genes could explain some of the mechanisms behind papillary stone-associated injury, we next asked if their urinary secretion can also screen for stone disease activity. To this end, we assayed for MMP7 and MMP9 in urine sample from a cohort of 55 patients with normal kidney function separated into 3 groups (**Figure 7**): 1) healthy controls with no known clinical history of stones, 2) inactive CaOx stone formers with a known clinical history of stones but without a recent stone event and 3) active CaOx stone formers undergoing surgery for stone removal. Our results show that the levels of MMP7 are increased in the urine of CaOx stone patients without active disease compared to healthy subjects (4.8±6.1 vs. 1.6 ng/mg Cr, respectively; p=0.01). MMP9 was also significantly higher in the urine of inactive stone patients compared to healthy controls (1.8±3.7 vs. 0.18±0.17 ng/mg Cr, respectively; p=0.02). In active stone formers, we observed even higher levels of urinary MMP7 (8.1±6.6 ng/mg Cr; p=0.0004 vs. controls and p=0.47 vs. inactive stone formers) and MMP9 (8.0±10.7 ng/mg Cr; p <0.01 p<0.0001 vs. controls and p=0.01 vs. inactive stone formers). These results suggest that both MMP7 and MMP9 are potentially important markers to monitor stone disease activity.

**Figure 7.**
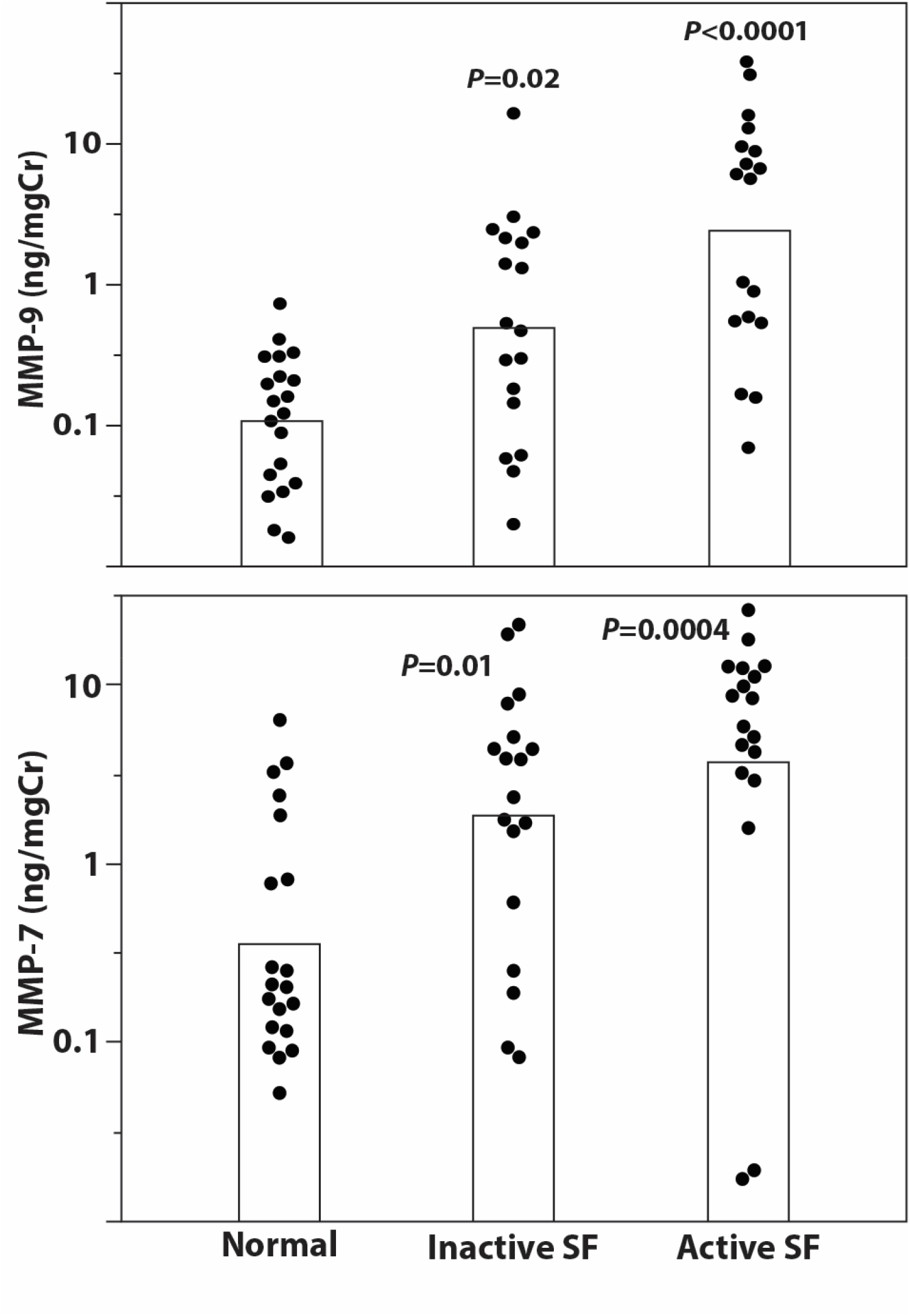
MMP7 and MMP9 levels are increased in urine of CaOx stone patients and correlate with disease activity. Urine samples were taken either from CaOx stone patients undergoing surgery for stone removal (active stone formers, SF) or from patients who had previously been stone formers (Inactive SF) or from healthy volunteers. Demographics and relevant clinical variables for each group are presented in Supplemental Table 2. Samples were assayed for urine (**A**) MMP7, (**B**) MMP9 and urine creatinine (Cr) by ELISA according to manufacturer’s instructions. Samples are plotted as log_10_(ng MMP/mg Cr).

## Discussion

In this work, we utilized snRNA sequencing, spatial transcriptomics, and large-scale multiplexed imaging to establish a spatially anchored cellular and molecular atlas of the renal papilla in non-stone and kidney stone disease tissues. A full landscape of papillary cells was defined, including the presence of papillary surface epithelial cells, stromal and immune cells, unique subtypes of principal cells, and an undifferentiated epithelial cell type which localized to regions of injury or mineral deposition. Despite the focal nature of mineral deposition in stone disease, we showed that injury pathways are globally upregulated across multiple cell types within the papilla. Commonly enriched signaling pathways in stone disease such as leukocyte (myeloid) immune activation, oxidative stress and matrix remodeling were demonstrated using orthogonal approaches, thereby enhancing our confidence in the findings. We defined the microenvironment of plaque as an active immune zone with antigen presenting inflammatory macrophages interacting with T cells, but also demonstrated the presence of an immune lifespan around mineral deposition ranging from inflammation to fibrosis. Finally, MMP7 and MMP9 were identified as two proteins linked to active stone disease and mineralization within the papilla. The levels of MMP9 and MMP7 in the urine were significantly higher in patients with history of stone disease compared to healthy controls, and correlated with disease activity.

Defining the cellular and molecular landscape in the human papilla is an important first step to understand the pathogenesis of nephrolithiasis and identify potential targets for therapy. An important strength of our study is the utilization of rare and highly valuable papillary biopsy specimens obtained from stone patients. The uncovered atlas of the renal papilla harbored expected cell types such as the principal, intercalated, ascending and descending thin limbs, and endothelial cells. Compared to cortico-medullary cells, papillary collecting duct cells displayed a unique gene expression signature that is consistent with the physiological milieu within the papilla. For example, the high expression of urea transporters is consistent with gene expression data from rodents (Fenton et al. 2005). Interestingly, the papilla was enriched in two types of principal cells, PC1 and PC2, which frequently co-localize in the same collecting ducts. PC2 exhibited a transcriptomic signature of cell stress and was more prevalent in specimens from subjects with stone disease. The biological significance of these two populations requires further investigation, particularly in establishing whether the injury signature of PC2 is a key contributor to the immune activation observed in stone disease.

Another cell type of interest was the “undifferentiated” snRNAseq epithelial cell cluster. Its signature did not align completely with a specific epithelial cell type but exhibited gene expression features of injury and degeneration. We posit that that this cluster may represent a final common injury phenotype derived from epithelial cells with multiple origins. This cell type was more frequently mapped in the stone samples and was associated with areas of mineralization in spatial transcriptomics (ST). In an orthogonal approach with CODEX, we uncovered an undifferentiated cell population with protein signatures overlapping with the transcriptomic features of the snRNAseq cluster. Spatial mapping suggested that these cells are likely a mixture of injured thin limbs and papillary epithelium, which is consistent with the location of these cells in the UMAP space. A signature of injury in thin limbs is concordant with previous data from our group showing that Randall’s plaque begins in thin limb cells (Evan et al. 2003). This injured population and its potential association with pathology in the papilla needs further exploration.

Comparing stone to reference samples using snRNA and spatial transcriptomics independently uncovered common DEGs and enriched pathways in stone disease. In snRNA and ST analyses, many of the genes and injury pathways uncovered were common among various cell types (epithelia, stromal and immune), suggesting that the injury signature associated with stone disease could reflect a global injury to the stone forming papilla. Our data provide important human tissue context to the previously reported hypotheses, and some of the experimental models of stone formation which involved inflammation, leukocyte activation (macrophage activity), ROS and ossification-like events (Khan 2012) (Khan et al. 2014) (Joshi, Wang, et al. 2015) (Joshi, Clapp, et al. 2015) (Taguchi et al. 2016; Taguchi et al. 2017). Our spatial transcriptomic mapping of gene expression in stone disease agrees with, and extends the work of Taguchi and colleagues, who explored genome-wide analysis of gene expression on renal papillary Randall’s Plaques (RP) and non-RP, and showed upregulation of *LCSN2, IL11*, and *PTGS1* in the RP patient tissue (Taguchi et al. 2017). RP has been previously described as the interstitial mineral deposition at the tip of the renal papillae that can serve as the origin for CaOx stone growth (Evan et al. 2006) (Daudon, Bazin, and Letavernier 2015). This immune active state in the regions of papillary mineralization has a molecular profile comparable to vascular inflammation leading to atherosclerotic disease, which has been proposed to play a pathogenic role in mineral deposition and stone disease (Kumada et al. 2004) (Abbas et al. 2014) (Bird and Khan 2017) (Li et al. 2020).

Our work particularly underscores the importance of immune system in the pathogenesis of stone disease, mapping significant populations of both the inflammatory M1 macrophage and the alternatively activated M2 macrophage within the papilla. An intermediate phenotype M1/M2 was particularly abundant in areas around mineral deposition, highlighting that these two phenotypes likely represent a spectrum that may modulate disease activity (Yunna et al. 2020) (Taguchi et al. 2021a; Taguchi et al. 2021b). This is consistent with previous reports by Khan and others from experimental models (de Water et al. 1999) (Khan 2004) (Joshi, Wang, et al. 2015; Taguchi et al. 2016) (Xi et al. 2019) (Taguchi et al. 2021a), but is uniquely demonstrated here in the human papilla. The presence of inflammatory macrophages and antigen presenting cells, surrounding by CD4+ T cells suggest a nidus for antigen presentation around plaque. Interestingly, in the same papillary specimen, we also uncovered a significant population of fibroblasts around an area of RP with less immune activity. This finding agrees with the matrix remodeling transcriptomic signature observed, which frequently results from chronic non-resolving inflammation. This is also consistent with recent findings by Canela et al. showing fibrosis and immune signature derived from imaging of CaOx stones with RP (Canela et al. 2021).

Cumulatively, these findings indicate that mineral deposits likely trigger a canonical immune injury pattern, which extends beyond the areas of mineralization and affect multiple cell types. Our data suggest that there could be various stages and biological responses to mineral deposition, ranging from an active acute inflammatory reaction, to established fibrosis secondary to chronic inflammation. Injury and inflammatory macrophage infiltration were demonstrated in biological replicates using 3D imaging and cytometry. Heavy mineral deposition was not obvious in the tissues analyzed by 3D imaging which raises a possibility of factors other than heavy mineral deposition leading to injury and immune activation, or could reflect earlier stages of diffuse papillary injury that precedes plaque deposition. Our data suggest that papillary injury triggers pro-fibrotic signaling and a vigorous matrix remodeling program. Indeed, our studies uncovered two matrix metalloproteinase molecules, MMP7 and MMP9, that are associated with stone disease and mineral deposition, respectively.

Matrix metalloproteinases (MMPs) degrade extracellular matrix proteins during growth, tissue remodeling and disease processes, and are secreted by various cell types including fibroblasts and leukocytes (including macrophages) (Cui, Hu, and Khalil 2017). Some MMPs control leukocyte migration and can modulate the immune response by biochemical cleavage of cytokines and chemokines (Elkington, Green, and Friedland 2009). MMP7, or matrilysin, is thought to modulate innate immunity and leukocyte influx, and plays a critical role in extracellular remodeling (Manicone and McGuire 2008) (Bulow and Boor 2019) (Burke 2004).

However, it is typically not expressed at the protein level in the renal cortex in healthy states (Zhou et al. 2017) (Fu et al. 2019) but its expression and activity are increased in the setting of kidney disease. There are no previous studies that have investigated MMP7 in the human papilla. Furthermore, there are conflicting data about the role of MMP7 in experimental models of kidney disease. MMP7 is thought to mediate kidney fibrosis in unilateral ureteral obstruction models, and its levels are elevated in patients with chronic kidney disease (CKD) (Zhou et al. 2017) (Surendran et al. 2004). However, MMP7 can also protect against acute kidney injury. In our papillary, non-stone, reference tissue, the transcriptomic signature of *MMP7* was predominantly localized to collecting duct cells. In stone biopsy specimens, the expression of *MMP7* was no longer limited to the collecting duct cells. snRNA and spatial transcriptomics were consistent in showing upregulation of *MMP7* expression in various cell types within the papilla. We propose that induction and release of MMP7 in various cell types is likely part of the matrix remodeling program induced by injury. The induction of MMP7 could therefore be a valuable indicator of papillary injury associated with stone disease.

MMP9 (macrophage gelatinase) is another MMP that was uncovered by our studies in association with mineral deposition. Our spatial transcriptomic analysis showed enriched expression of *MMP9* in areas of mineral deposition, which co-localizes with osteopontin (*SSP1*) expression. Indeed, *MMP9* gene polymorphisms are associated with nephrolithiasis (Mehde et al. 2018). *MMP9* is upregulated by classically activated macrophages during an inflammatory response, and is also expressed by other inflammatory cells and osteoclasts (Hanania et al. 2012) (Wang et al. 2013) (Jager et al. 2016). MMP9 is proposed to play a role in renal fibrosis and epithelial to mesenchymal transition (Tan et al. 2010) (Tan et al. 2013). It also interacts with osteopontin to enhance macrophage chemotaxis and fibrosis (Tan et al. 2013). More recently, experimental studies by Wu et. al. suggested that activation of ROS in kidney tubular epithelial cells by way of the NK-κB/MMP-9 pathway promotes crystal deposition in the kidney (Wu et al. 2021). Therefore, upregulation of MMP9 could be an indicator of crystal deposition, immune activation, and transition towards a profibrotic phenotype.

Since our tissue work suggested a potential role for MMP7 and MMP9, we investigated whether measuring these molecules in the urine could be useful in patients with stone disease. We found that both MMP7 and MMP9 are increased in patients with stone disease and their levels correlate with disease activity. MMP7 has been previously studied as a potential biomarker for predicting risk of acute kidney injury and CKD progression (Liu, Tan, and Liu 2020). MMP7 correlated with fibrosis scores on kidney biopsies and was also found useful to predict IgA nephropathy progression (Yang et al. 2020). More recently, MMP7 levels were also elevated in patients with hypertension who developed CKD (Sarangi et al. 2022). MMP9 levels have also been studied in the setting of diabetic kidney disease and found to be elevated in patients with macroalbuminuria (Pulido-Olmo et al. 2016) (Garcia-Fernandez et al. 2020). To our knowledge, our study is the first to show that the urine levels of MMP7 and MMP9 are elevated in patients with stone disease compared to non-stone forming kidney subjects, particularly in the absence of a clinically detectable kidney function impairment. MMP7 and MMP9 urine levels are higher, particularly for MMP9, in patients with symptomatic active stone disease. Large studies are needed to define the utility of these markers in the management of patients with kidney stone disease. For example, longitudinal measurements of these markers and their changes may help predict stone recurrence and response to therapy, supplementing computed tomography to reduce lifetime radiation exposure in stone formers. Indeed, our results provide a solid rationale to study these markers in the context of a large clinical trial.

Our study has limitations. Renal papilla samples from stone formers are challenging to obtain as they require a biopsy during a nephrolithotomy surgical procedure. Fortunately, our investigative team has substantial experience obtaining these samples (Evan et al. 2006) (Evan et al. 2018). The number of tissue specimens used to generate the transcriptomic and imaging datasets was relatively small, underscoring the importance and scarcity of these samples. Despite the sample size, the findings were validated by orthogonal methods and the importance of the pathways and molecules uncovered were ultimately linked to urinary measurements in a larger clinical cohort. Further, the co-clustering of nuclei from the renal papilla with a publicly available snRNAseq whole kidney atlas (of more than 200,000 nuclei) allowed substantially more power to distinguish granular cell subtypes and cell states. Due to the cost prohibitive nature of multi-omics studies, our approach of discovery in a small sample size and validation in a larger cohort serves as a viable model for hypothesis driven future studies economizing on tissue and maximizing validation efforts on samples collected through non-invasive methods over time. As discussed, the findings of elevated MMPs could also be observed in other forms of kidney disease. However, the cohorts used here had normal kidney function, and the potential utility of these markers in tracking stone disease activity, particularly in patients with normal kidney function, is thereby not diminished. We studied CaOx kidney stone formers, which is the most common type of kidney stones. The applicability of our findings to other types of stone disease will need to be established by additional studies.

In conclusion, our integrated multi-omics investigation uncovered the complexity of the human kidney papilla and provides important insights into the role of immune system activation and mineral deposition. We established the diffuse injury associated with nephrolithiasis and highlight the importance of a matrix remodeling program in the kidney papilla. Particularly, we identified MMP7 and MMP9 as potential molecules that may serve as noninvasive markers of kidney stone disease course and activity. This investigation provides a strong foundation for future studies that explore the pathogenesis of kidney stone disease and the application of precision medicine to patients with nephrolithiasis.

## Acknowledgements

We thank Sharon Bledsoe, Jennifer Stashevsky, and Katrina Kelly for help with tissue specimens. We also thank KPMP participants and the HuBMAP consortium.

