## Supplemental Figures for "A spatially anchored transcriptomic atlas of the human kidney papilla identifies significant immune injury and matrix remodeling in patients with stone disease"

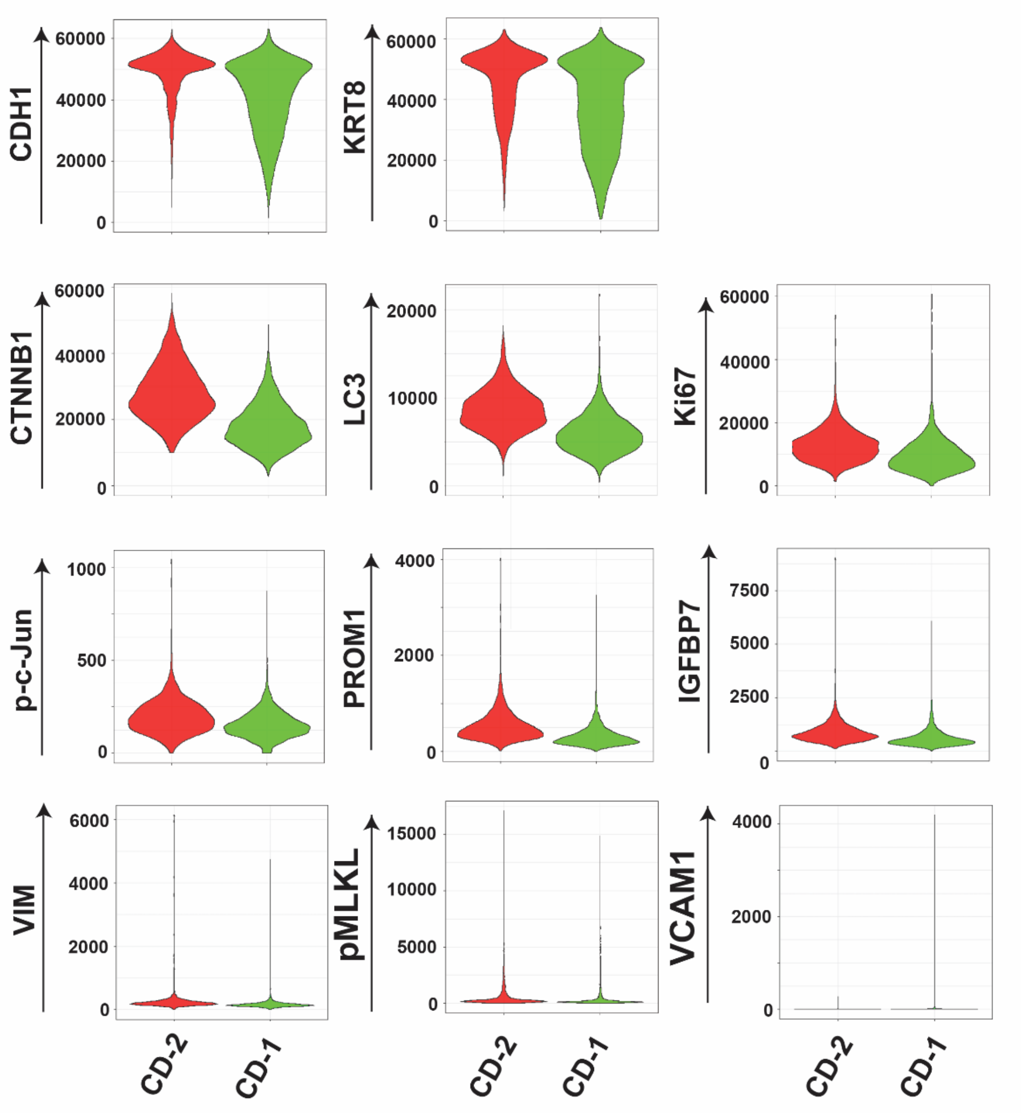


**Supplemental Figure 1. Injury profile of CD-1 and CD-2.** Violin plots show the mean fluorescence intensity distribution levels of cell and injury markers for two populations of collecting duct cells, CD-1 and CD-2, detected in reference papilla by CODEX multiplex imaging. The two populations had comparable distributions of the collecting duct markers, CDH1 and KRT8, but CD-2 had higher expression for injury markers such CTNNB1 (β-catenin), LC3, Ki67, p-c-Jun, PROM1, pMLKL and IGFBP7.

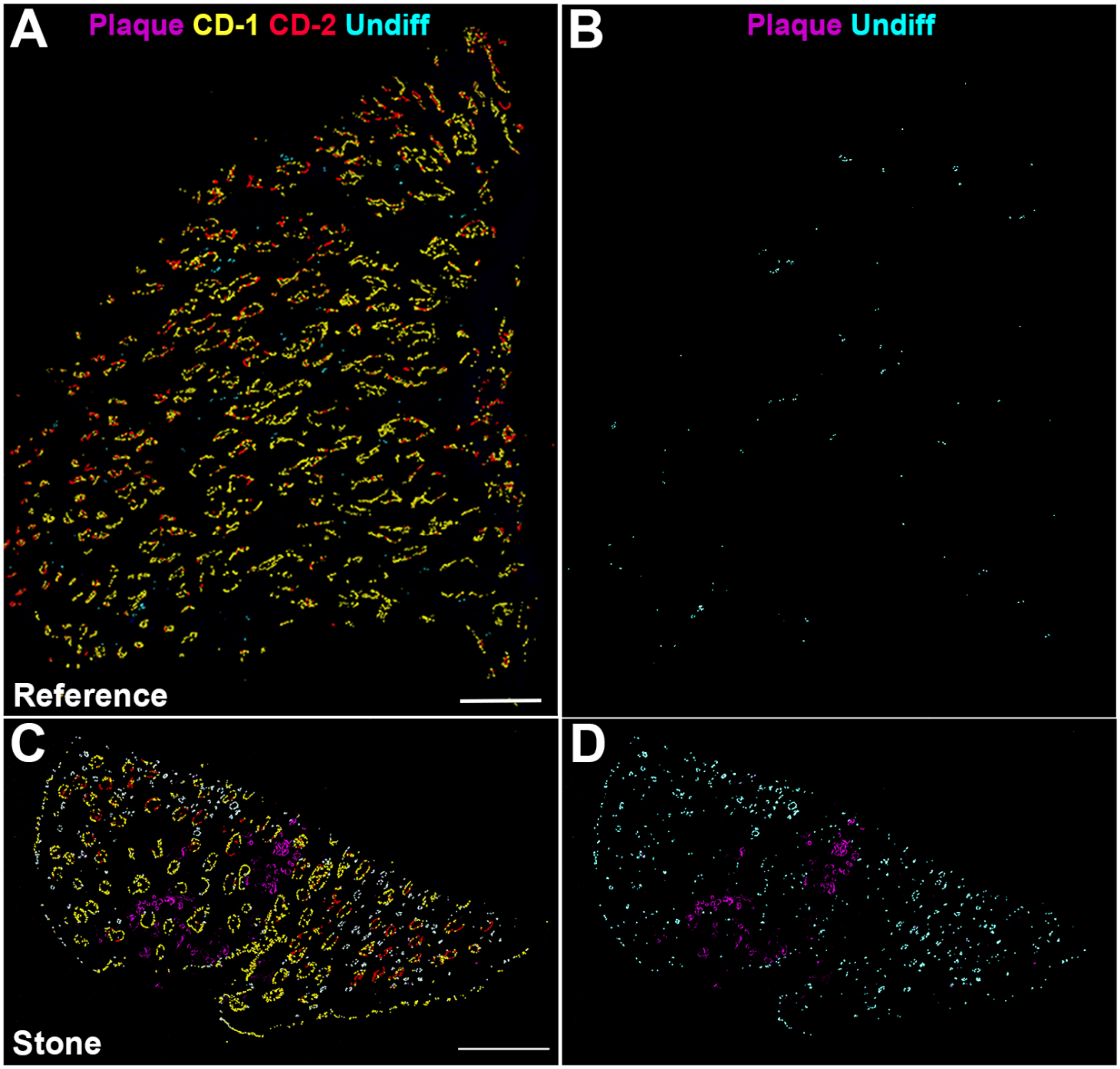


**Supplemental Figure 2. CODEX imaging of collecting duct and undifferentiated epithelial cell clusters in healthy and stone specimens.** Reference (A,B) and Stone (C,D) show various degree of distribution of CD-1 and CD-2 cells. Mineral deposition or Randall's plaque is autofluorescent and is clearly visible in the stone specimen (C and D). Undifferentiated cells (Undiff) are sparsely in reference tissue (B) but are diffusely abundant in the stone specimen (D). Scale bars = 0.5 mm.

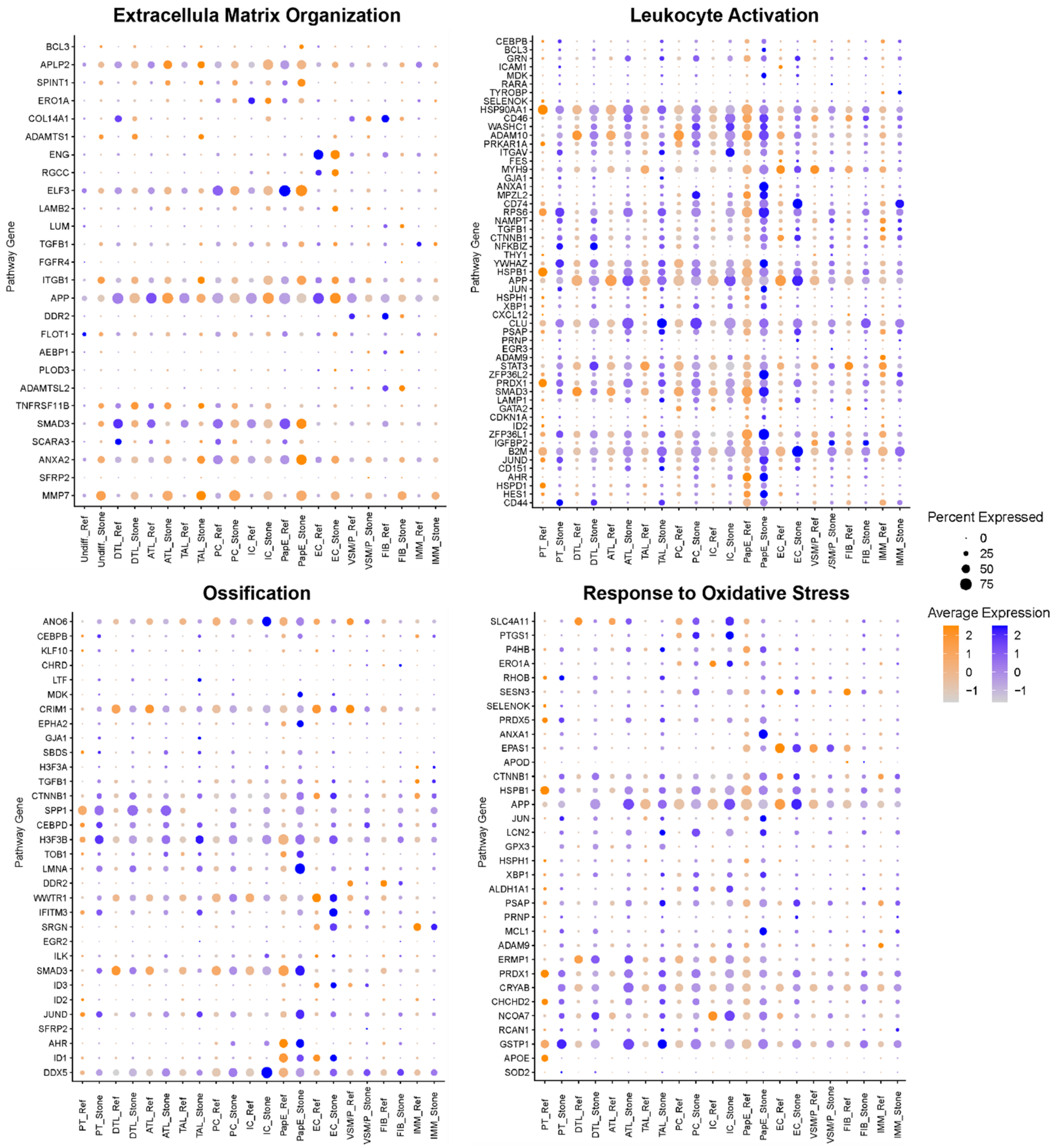


**Supplemental Figure 3. Cell type expression of select genes from pathways enriched in stone versus reference patient papilla biopsies.** Genes in relevant pathways (Extracellular Matrix Organization-GO:0030198, Leukoctye Activation-GO:0045321, Ossification-GO:0001503 and Response to Oxidative Stress: GO:0006979) that were significantly increased in all stone biopsies as detected by snRNAseq analysis were compared by dot plot across cell types identified based on their transcriptomic signatures relative to the snRNAseq atlas. Undiff. = undifferentiated; DTL = descending thin limb; ATL = ascending thin limb; TAL = thick ascending limb; PC = principal cell; IC = intercalated cell; PapE = papillary epithelium; EC = epithelial cell; VSM/P = vascular smooth muscle cell; FIB = fibroblast; IMM = immune cell.

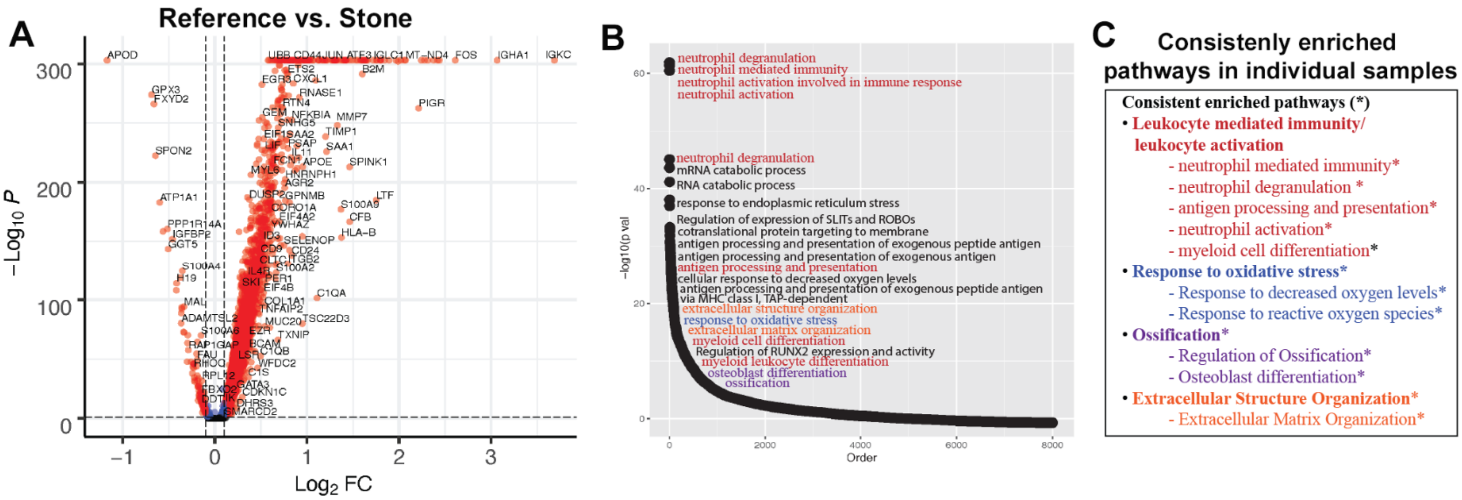


**Supplemental Figure 4: Differentially expressed genes and pathway in stone papillary specimens using pseudo-bulk analysis from spatial transcriptomics** (A) Volcano plot displaying differentially expressed genes (DEGs) between reference (left) and CaOx stone samples. (B) Differentially enriched pathways associated with DEGs from transcriptomic comparisons are illustrated in the pathway curve. Consistently enriched pathways that were common in each individual sample (N=3) compared to reference are listed in (C).

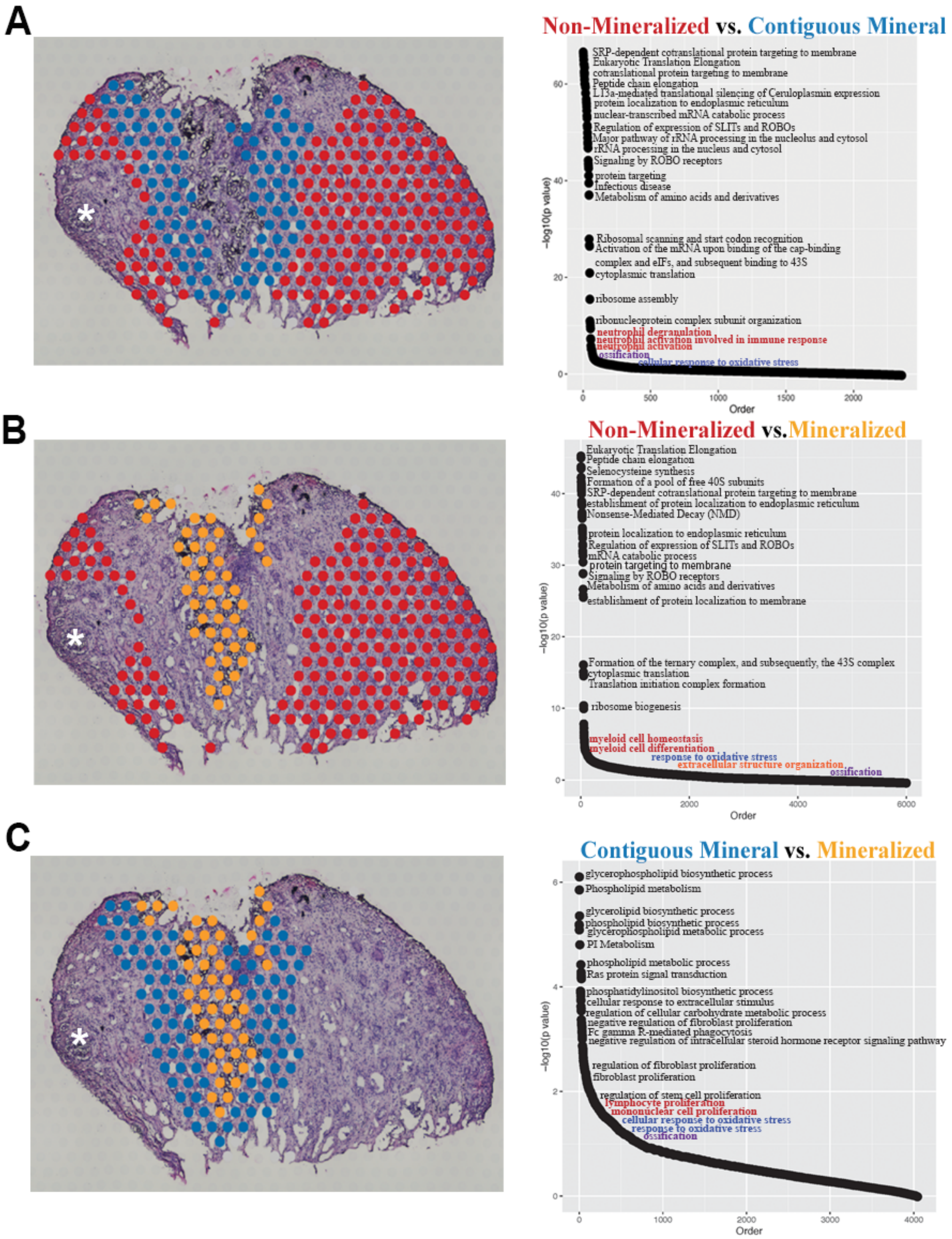


**Supplemental Figure 5: Differentially enriched pathways based on regional analysis and spatial association with mineralization.** Pathways linked to myeloid activation, oxidative stress, matrix remodeling and ossification are colored in red, blue, orangs and purple, respectively.
