## Supplemental tables for "A spatially anchored transcriptomic atlas of the human kidney papilla identifies significant immune injury and matrix remodeling in patients with stone disease"

**Supplemental Table 1: Papilla biopsy usage in spatial assays and post-sequencing quality control**

| Papilla Sample | Age | Sex | Tissue type | 3D imaging | CODEX | ST | ST Quality Control | | |
|---|---|---|---|---|---|---|---|---|---|
| | | | | | | | **Number of spots** | **Mapping to exons** (%) | **Mapping under tissue** (%) |
| **1** | 39 | F | Ref | X | | X | 2438 | 0.92 | 0.83 |
| **2** | 50 | M | Ref | X | X | | | | |
| **3** | 59 | F | Ref | X | | | | | |
| **4** | 63 | F | Ref | X | | | | | |
| **5** | 53 | F | CaOx | X | X | X | 817 | 0.53 | 0.53 |
| **6** | 44 | M | CaOx | X | | | | | |
| **7** | 36 | M | CaOx | X | | X | 1085 | 0.82 | 0.79 |
| **8** | 62 | F | CaOx | | | X | 939 | 0.86 | 0.75 |
| **9** | 63 | F | CaOx | X | | | | | |

ST = spatial transcriptomics

**Supplemental Table 2: Clinical summary of urine donors for MMP7/9 studies.**

| | **Normal** N= 20 | **Non-Active SF** N= 18 | **Active SF** N= 18 | ***P*** |
|---|---|---|---|---|
| **Age** | 41 +/- 4.2 | 39 +/- 4.5 | 47 +/- 5.2 | ns |
| **Sex** (% male) | 50 | 67 | 40 | ns |
| **Race** (% white, non-hispanic) | 100 | 100 | 100 | ns |
| **EGFR** (ml/min) | 93.2 +/- 9.2 | 98.10 +/- 9.4 | 88.9 +/- 11.7 | ns |
| **Serum Creatinine** (mg/dl) | 0.92 +/- 0.04 | 0.93 +/- 0.06 | 1.00 +/- 0.23 | ns |
| **Urine Creatinine** (mg/dl) | 152.3 +/- 30.0 | 119.9 +/- 28.3 | 121.3 +/- 37.2 | ns |
| **Diabetes** (%) | 0 | 0 | 5.6 | N/A |
| **HTN** (%) | 0 | 0 | 0 | N/A |
| **Cardiac Disease** (%) | 0 | 0 | 0 | N/A |

EGFR= Estimated glomerular filtration rate

**Supplemental Table 3: antibodies used in CODEX assay**

| Antibody | Significance | Clone | Supplier | Catalog Number |
|---|---|---|---|---|
| Ki67 | Proliferating cells | B56 | Akoya | 4250019 |
| CD3 | Pan T cells | UCHT1 | Akoya | 4350008 |
| CD4 | CD4+ T cells | SK3 | Akoya | 4350010 |
| CD8 | CD8+ t cells | SK1 | Akoya | 4150004 |
| CD11c | resident dendritic cells | S-HCL-3 | Akoya | 4350012 |
| CD31 | endothelial cells | WM59 | Akoya | 4250009 |
| CD20 | B cells | L26 | Akoya | 4150018 |
| CD45 | pan leukocyte markers | HI30 | Akoya | 4150003 |
| CD45RO | memory T cells | UCHL1 | Akoya | 4250023 |
| HLA-DR | antigen presenter cells | L243 | Akoya | 4250006 |
| CD90 | PT, fibroblasts, activated endothelial cells | SE10 | Akoya | 4150021 |
| E-cadherin | DCT, CD, loop of henle | 4A2C7 | Akoya | 4250021 |
| b-catenin | Tubular epethelium | 12F7 | Akoya | 4450036 |
| Cytokeratin8 | CNT and CD | TS1 | NovusBio | NBP2-34501-0.1mg |
| Uromodulin | TAL | Polyclonal | R&D | AF5144 |
| a-sma | myofibroblast, arterioles | 1A4 | Invitrogen | 14-9760-82 |
| PROM1 (CD133) | fibrosis | AC133 | Miltenyi Biotec | 130-090-422 |
| MPO | neutrophils | Polyclonal | Abcam | ab9535 |
| CD68 | activated macrophages | KP1 | ThermoFisher | 14-0688-82 |
| IGFBP7 | injury | Polyclonal | Acris/Origene | AP01109PU-S |
| p- c-Jun | stress kinase pathway | D47G9 | Cell Signalling | 3270BF |
| CD206 | M2 | Polyclonal | R&D | AF2534 |
| OPN | SSP1/osteopontin | AKm2A1 | Santa Cruz | sc21742 |
| ERG | Endothelial Nuclei | EPR3864 | Abcam | ab92513 |
| AQP1 | PT, TDL | 1/22 | Santa Cruz | sc-32737-X |

| | | | | |
|---|---|---|---|---|
| Citruline H3 | netosis | 7C10 | Acris/Origene | AM10179PU-N |
| Vimentin | Fibroblasts | RV202 | BD Pharmingen | 550513 |
| FOXP3 | injury | 236A/E7 | Thermo Fisher | 14-4777-82 |
| VCAM1 | non-reparining epi cells | EPR5047 | Abcam | ab271899 |
| phosphoMLKL | necroptosis | D6H3V (S358) | Cell Signaling | 91689BF |
| Fibronectin | Injury, pre-collagen | F1 | Abcam | ab271831 |
| LC3 | autophagy | Polyclonal | Sigma Aldrich | L8918-25UL |

Antibodies purchased from Akoya were conjugated by vendor. Antibodies from other vendors were conjugated in-house using Akoya conjugation kits as described in methods
